# Magnetic Nanoprobes for Spatio-Mechanical Manipulation in Single Cells

**DOI:** 10.1101/2021.08.16.455233

**Authors:** Iuliia P. Novoselova, Andreas Neusch, Julia-Sarita Brand, Marius Otten, Mohammad Reza Safari, Nina Bartels, Matthias Karg, Michael Farle, Ulf Wiedwald, Cornelia Monzel

**Author notes:** These authors contributed equally to this work (Iuliia P. Novoselova, Andreas Neusch).

## Abstract

Magnetic nanoparticles (MNPs) are widely known as valuable agents for biomedical applications. Yet, for their successful application within cells they need to fulfill a variety of demands such as monodispersity, biocompatibility or sufficient magnetic response. Given these prerequisites, MNPs may be used for remote, non-invasive manipulation, where their spatial redistribution or force response in a magnetic field provides a fine-tunable stimulus to a cell. Here, we investigate the properties of two different MNPs and their suitability for spatio-mechanical manipulations: semisynthetic magnetoferritin nanoparticles and fully synthetic ‘nanoflower’-shaped iron-oxide nanoparticles. Next to characterizing their structure, surface potential and magnetic response, we monitor the MNP performance in a living cell environment using fluorescence microscopy and confirm their biocompatibility. We then demonstrate their capability to spatially redistribute and to respond to magnetic force gradients inside a cell. Our remote manipulation assays present these tailored magnetic materials as suitable agents for applications in magnetogenetics, biomedicine or nanomaterial research.

## 1. Introduction

Functional magnetic nanoscale particles (MNPs) are widely employed in biotechnology and nanomedicine to study fundamental biological processes as well as to develop enhanced diagnostic and treatment strategies, the most prominent examples being smart drug delivery, contrast enhancement in imaging, magnetic separation of molecules, or magnetic particle hyperthermia [1]. In addition, for subcellular applications in fundamental studies, their manipulation via magnetic tweezers demonstrated to be beneficial for the study of organelles, proteins, and biomolecules within the cell environment [2], [3].

Iron oxide nanoparticles (NPs) are due to their versatile applicability and biocompatibility one of the most popular compounds for biomagnetics studies [4]–[7]. Interest in iron oxide NPs rises because they can be synthesized in various shapes, sizes and in large amounts (e.g. via thermal decomposition or laser target evaporation) enabling a cost-effective production. Previous studies demonstrated successful NP delivery into living tissues, their spatial manipulation by external magnetic fields [8], [9], as well as their controlled heating [10]. Recent studies, however, indicate that these nanoprobes need to reconcile a variety of demands which are difficult to achieve simultaneously [1], [11]. First, NPs need to be biocompatible. Second, they need to be sufficiently monodisperse in terms of size, surface charge and functional motifs in order to respond similarly to an external magnetic stimulus (DC or AC field). Finally, they need to be functionalized to specifically target particular biological sites as well as to make the particle chemically inert. Here, we characterize a new class of ‘nanoflower’-shaped multicore iron oxide NPs, called synomag, and discuss possible applications.

Semisynthetic particles are another new class of NPs which can be tailored to meet the listed demands. Highly successful examples are NPs made of proteins belonging to ferritins, a group of proteins which naturally store iron in an organism to maintain its iron homeostasis [12], [13]. For example, human ferritin (Ft) is a globular protein cage of this family, consisting of 24 subunits, 12 heavy and 12 light chains. Interestingly, the heavy-chain-ferritin (HCF) subunit exhibits ferroxidase activity [13], [14] which can be exploited in Fenton-like reactions to synthesize a magnetic crystal into the cage. Previously, it was shown that a superparamagnetic iron oxide core can be grown within the cage [15]. This is particularly beneficial for subcellular applications where particle chain formation due to magnetic dipole-dipole interactions should be prevented. Moreover, due to the unique protein structure, these NP templates are biocompatible by nature and exhibit a well-defined size with a strict upper size limit along with shape uniformity [16]. In addition, the protein cage can serve as a scaffold for bio-orthogonal tagging, and via genetically encodable tags, precise control over the tag number is achieved. Such genetically modified human ferritin, henceforth termed ‘magnetoferritin’ (MFt) is the second MNP presented in this work.

If a MNP exhibits the desired superparamagnetic, biofunctional and biocompatible properties, it is imperative to evaluate the MNP in conjunction with an applied magnetic field. Any spatial, mechanical or thermal stimulus to be transferred to a biological entity of interest will heavily depend on both, the NP magnetic responses and the externally applied field. For example, for localized MNP heating, the dissipated heat is proportional to the AC susceptibility of the NP and the square of the magnetizing field [17]. In case of force applications, the product of the NP’s magnetic moment and the magnetic field gradient have to be known. The localized heat or applied force may then be used to switch molecular activity states and to eventually influence a cellular function. Such remote and finely tunable magnetic manipulation approach is commonly referred to as Magnetogenetics. Several recent studies demonstrate how these stimuli (heat, force or spatial redistribution) enable active probing of fundamental cell signaling functions, such as action potential formation in neuronal cells [18], cell signaling for differentiation or migration [9], [19], or tumorigenesis [20]. Hence, Magnetogenetics can provide a rich toolkit to study fundamental processes in individual cells. An overview of intriguing biological and medical questions is reviewed in Monzel et al. [21] and Pankhurst et al. [17] along with information about the required magnetic fields and gradients.

## 2. Materials and Methods

### Ferritin Cage Expression and Purification

monomeric enhanced green fluorescent protein (mEGFP) tagged heavy chain ferritin (HCF) plasmid (mEGFP::HCF) was a kind gift from the Coppey/Hajj Lab at Laboratoire Physico-Chimie, Institut Curie, Paris, France and Piehler Lab at University Osnabrück, Germany. For site-specific targeting of proteins, mEGFP was fused to the N-terminus of HCF containing 6 amino acids as linker by cassette cloning as described before [19]. For bacterial expression of mEGFP::HCF in *E.coli* BL21-CodonPlus (DE3)-RIPL Competent Cells (Agilent Technologies, Santa Clara, CA, USA), the cDNA of the fusion protein was cloned in pET21a (Merck KGaA, Darmstadt, Germany). For preparative overexpression the *E. coli* BL21 were transformed with the plasmid and the culture was grown from a single colony in 2xYT medium at 37 °C up to an OD_600_ of 0.6 to 0.8. Plasmid expression was induced with 1 mM IPTG (Sigma-Aldrich, St. Louis, MO, USA) and the culture was grown further at 16 °C overnight. Harvested cells were washed in Phosphate Buffered Saline (PBS) and finally resuspended in HEPES buffer (150 mM NaCl, 50 mM HEPES, pH 8.0). The obtained cell suspension was treated with Protease Inhibitor Cocktail and PMSF to protect the protein from degradation during the following cell disruption via a homogenizer (M110P Microfluidizer, Microfluidics, Westwood, MA, USA). After centrifugation, the supernatant containing mEGFP::HCF was purified by heat shock (70 °C, 15 min). Subsequently, mEGFP-tagged ferritin particles were cleaned up using ammonium sulfate precipitation, first at 200 g/l, then at 300 g/l, to precipitate proteins of different solubility than the desired protein. After the last step, the pellet containing ferritin was resuspended and dialyzed overnight in HEPES buffer (20 mM HEPES, 100 mM NaCl, pH 8.0). The resulting sample was loaded onto a size exclusion column (HiPrep 16/60 Sephacryl S400 High Resolution, GE Healthcare, Chicago, IL, USA) equilibrated with filtered (0.2 μm) and degassed buffer (20 mM HEPES, 100 mM NaCl, pH 8.0). All chromatography steps were performed in a FPLC system (Äkta Explorer, GE Healthcare). 12% Sodium dodecyl sulphate–polyacrylamide gel electrophoresis (SDS-PAGE) was used to confirm the purity of the final product as well as the effectiveness of each individual purification step [22].

### Coupling of methoxy-PEG2000-NHS

methoxy-(polyethylene glycol)-N-hydroxysuccinimid (Molecular Weight (MW): 2000 Da) was purchased from Sigma-Aldrich. 30 μl of 100 mM methoxy-PEG2000-NHS in dry DMSO were added to 100 μl mEGFP::HCF (10 μM) in 20 mM HEPES, 100 mM NaCl, pH 8.0. The reaction mixture was incubated for 2 h at room temperature on a rotator. PEGylated mEGFP tagged ferritin was purified with PD10 desalting columns (GE Healthcare) equilibrated with 20mM HEPES, 100mM NaCl at pH 8.0. The integrity and physicochemical properties of PEGylated mEGFP::HCF were examined by SDS-PAGE. The protein concentration was determined by UV-Vis absorption spectroscopy (NanoDrop 2000, Thermo Fisher Scientific Inc., Waltham, MA, USA).

### Iron Oxide Core Synthesis in Ferritin Cages

purified PEGylated ferritin cages were used to synthesize a magnetite core inside the cage using 1 mg/ml protein concentration diluted in 25 ml of 100 mM NaCl (Sigma-Aldrich). Solution of 30 mM ammonium iron (II) sulfate hexahydrate (Sigma-Aldrich) was used as an iron source and added to the reaction vessel at a constant rate of 211 μl/min at an excess of about 5000 Fe atoms per ferritin cage. Next to the iron source, hydrogen peroxide H_2_O_2_ of 5 mM was simultaneously syringe-pumped into the vessel at the same rate. During the synthesis, the reaction vessel was kept at 65 °C under positive N_2_ pressure and the pH was maintained dynamically at 8.5 with 100 mM NaOH by an automatic titrator (Titration Excellence T5, Mettler-Toledo, Columbus, OH, USA). During the oxidation process, the green-toned solution became evenly yellow-brown. Once magnetic loading was done, 200 μl of 300 mM sodium citrate was added to chelate any free iron. Synthesized MNPs were treated by centrifugation at 19.000 g for 30 min at 4 °C and 0.2 μm PTFE filtering to remove potential iron oxide aggregates formed outside of the cages. The final product was stored at -80 °C to prevent protein degradation till further experiments were performed.

### Synomag NPs

‘nanoflower’-shaped iron oxide particles were commercially available from micromod GmbH (Rostock, Germany). Since some of the NPs exhibited free amine groups on their surface, coupling of PEG2000-COOH was performed to generate a passivation surface according to the protocol given above ‘Coupling of methoxy-PEG2000-NHS’.

### Transmission electron microscopy (TEM)

ferritin cages in buffer solution were dropped onto Formvar coated nickel grids (200 mesh, S162N-100, Plano GmbH, Wetzlar, Germany) and left to sediment for 1 min. Solution remaining on the grid was removed between each step using filter paper. After sedimentation, the sample was negatively stained twice for 3 - 5 s and 30 s by subsequently placing the grid onto two separate drops of 2 % uranyl acetate. The remaining solution was removed and the grid was left to dry on air.

Imaging was performed on a Jeol JEM-2100Plus (Akishima, Tokyo, Japan) operating in bright field mode at an acceleration voltage of 80 kV. The average size was obtained by statistical analysis using Fiji [23] of 275-347 NPs for synomag, stained ferritin shells, and unstained magnetic cores in ferritin NPs, respectively.

### Dynamic light scattering (DLS) and electrophoretic light scattering (ELS)

measurements were carried out on Zetasizer Nano ZS (Malvern Panalytical Ltd, Malvern, UK) equipped with a He-Ne laser working at a wavelength of 633 nm. Data was recorded using a detection angle of 173°. For DLS, 5 subsequent measurements containing 50 sub-runs were taken (each sub-run duration 10 s). Reported hydrodynamic diameters *D*_H_ refer to the peak value of the log-normal fit applied to the DLS number distributions. For ELS, stabilizing buffers showed a conductivity of about 10-15 mS/cm (measured conductivity of deionized water is below 0.1 mS/cm), thus, measurements were carried out at a reduced voltage of 40 V to protect measuring cell electrodes from deterioration. Reported results represent averages from 10 measurements, 30 sub-runs each along with standard deviations.

### Magnetometry

buffers from the samples were diluted to reduce the amount of salts. Liquid samples were dried using a rotational evaporator. Dried powders (~10 mg) were compacted into synthetic sample holders for vibrating sample magnetometry (VSM) in a PPMS DynaCool system (Quantum Design GmbH, Darmstadt, Germany). The magnetic response was measured up to 4 T at 5-300 K, and zero-field-cooling/field-warming sequences (ZFC/FW) were recorded at 5 mT.

### Cell Handling

HeLa wild type cells (ATCC^®^ CCL-2^™^, ATCC, Manassas, VA, USA) were cultivated at 37 °C, 5% CO_2_ in Dulbecco’s Modified Eagle Medium (DMEM) medium (Thermo Fisher Scientific) supplemented with 10% fetal calf serum (FCS) and 1% PenStrep (Thermo Fisher Scientific).

### Cell Viability Assay

to assess possible negative effects of MNPs on cell viability, a standard CellTiter-Blue (CTB) Assay from Promega (Fitchburg, WI, USA) was carried out for both empty ferritin cages and synomag samples. This assay is based on the conversion of resazurin to resorufin by living cells. This reduction causes a shift in the dye’s fluorescence. In a 96 well plate about 25.000 cells were seeded per well and grown in the incubator (37 °C; 5% CO_2_) overnight. Afterwards, cells were incubated with increasing amounts of MNPs. Untreated cells and cells incubated only with the MNP containing medium served as positive control. Incubation with MNPs was carried out for 24 h. As negative control, untreated cells were exposed to 0.1% Triton 30 min before CTB addition. After incubation, cells were washed with DPBS and incubated for 4 h in a mixture of DMEM and CTB (9:1). The emission of the supernatant at 590 nm was measured (excitation: 560 nm) in a plate reader (infinite M200 Pro, TECAN, Männedorf, Switzerland). The obtained signal is linearly proportional to the number of living cells inside the sample.

### Imaging, nanoparticle incubation and microinjection, image analysis

for imaging and manipulation, HeLa wild type (WT) cells were plated on sterilized glass-coverslips in 35 mm cell-culture dishes at about 50% confluency. Prior to experiments, cells were washed with a PBS buffer and re-incubated in preheated Leibovitz medium (L15, Thermo Fisher Scientific) supplemented with 10% fetal calf serum (FCS) and 1% PenStrep (Thermo Fisher Scientific). Imaging was performed in a heating chamber preheated to 37 °C and placed on an inverted microscope (IX83 from Olympus, Shinjuku, Tokyo, Japan) equipped with a 60x oil-objective with N.A. 1.25 and phase contrast Ph3. For fluorescent optical MNPs detection and microinjection, cells were seeded to adhere in 8-well rectangular chambers equipped with a glass bottom (Sarstedt, Nümbrecht, Germany) overnight in an incubator at 37 °C, 5% CO_2_. MNPs were centrifuged (19.000 g for 10 min) and filtered through PTFE filters with 0.2 μm cutoff (Filtropur S 0.2, Sarstedt). After this cleaning routine, cells were incubated for either 6 or 24 h with MNP concentrations of 0.1 - 1.0 mg/ml. Directly before microscoping imaging, cells were washed with preheated PBS and immediately incubated in L15 medium supplemented with FCS and PenStrep as before. MNPs were injected into cells using a micro manipulation system (Inject-Man 4 and FemtoJet 4i, Eppendorf, Hamburg, Germany) with microinjection capillaries (Femtotip II, inner diameter of 500 nm, Eppendorf). For injection, a concentration of 3 mg/ml for each MNP system was used and capillary pressures were set to a pressure difference between the needle tip and the environment of 10 to 20 hPa.

The MNP uptake and temporal change inside the cells was analyzed using an in-house developed analysis routine written in Matlab (R2019b, The Mathworks Inc., Natick, MA) and ImageJ (version 1.49v, NIH, USA). For their detection, phase contrast and fluorescence image recordings were used. Cells borders were detected using a canny edge detection algorithm. If necessary, cell borders were corrected manually. Single cells were then numbered and cellular fluorescence intensity was attributed to each corresponding cell. The intensity by area was determined by the quotient of fluorescence intensity over cell area for each cell.

### Magnetic Tip Configuration

the magnetic tip is constructed from two magnets (see also Figure 5; cube: NdFeB, 5 × 5 × 5 mm, gold-plated, product number W-05-G; cuboid: NdFeB, 10 × 4 × 1.2 mm, gold-plated, product number: Q-10-4-1.2-G; both from Webcraft GmbH, Gottmadingen, Germany) and a 0.4 mm diameter polished steel wire (product number 1416, from Röslau Stahldraht, Germany) attached to it. During the remote manipulation experiments, the magnetic tip was brought to the HeLa WT cell edge at a moderate distance of 10 - 20 μm for 10-15 min and removed thereafter.

Simulations of the magnetic tip stray field distribution were performed using COMSOL Multiphysics^®^ software (COMSOL Inc., Stockholm, Sweden) with an AC/DC electromagnetics module. The magnetization curve of a Neodymium magnet was implemented into the COMSOL library as material reference. According to the mentioned tip wire material, spring steel was chosen as material reference for the tip wire. This data was used to simulate the field distribution of the magnetic tip in the configuration used for remote manipulation experiments.

## 3. Results and Discussion

Magnetoferritins (MFts) and synomag are two structurally and bio-functionally complementary particles, which were chosen to test their suitability for the manipulation of biological functions inside cells. In case of MFts, the ferritin protein cages served as templates, which were further tailored to feature cytosolic stealth properties as well as to fluoresce for microscopic observation. To this end, we used a bacterial expression vector of the heavy chain subunit of the ferritin protein (HCF) with a mEGFP genetically fused to it. Note, that mEGFP is pointing away from the ferritin surface as any encapsulation of the fluorescent molecules inside the cage structure is sterically inhibited. In order to render ferritin cages chemically inert and mobile within the living cell environment, a polyethylene glycol (PEG) coupling to its surface with 2 kDa PEG was performed as previously described [19]. The PEGylation was suggested to not only result in cytosolic stealth properties but also to improve cage stability during the magnetic core synthesis at 65 °C [16]. We synthesize a pure magnetic iron oxide core into the functionalized cage by taking advantage of the intrinsic ferroxidase activity of the heavy chain ferritin subunit in a Fenton-like reaction. For details of the MFt synthesis refer to the Materials and Methods section of this work and Lisse et al. [19]. On the other hand, the fully synthetic core-shell MNPs synomag were used. These consist of iron oxide cores embedded into a dextran shell forming an inhomogeneous sphere-like morphology [24]. Due to their production via co-precipitation, synomag are produced at much higher quantities than MFt, albeit with a larger variety in size. Thus, in subsequent steps, their size distribution was narrowed via separation in high gradient magnetic fields. Synomag are commercially available with amine reactive surface groups for covalent coupling of a biofunctional molecule such as carboxyl-polyethylene glycol (COOH-PEG). While synomag MNPs previously proved to be highly suitable for magnetic hyperthermia [25] and magnetic particle imaging [26], they are probed herein with regard to their force and spatial manipulation capability.

To characterize both MNPs, we performed structural and morphological analyses of MFt and synomag particles whose schematic representations are shown in Figure 1(a) [27]. Dynamic Light scattering (DLS) was used to determine the hydrodynamic particle size and polydispersity. The data of both particles showed a narrow, monomodal size distribution (Figure 1(b)) from which the effective hydrodynamic diameter *D_H_* was extracted. The peak value of the number distribution revealed a *D_H_* of 39.1 ± 2.5 nm and 39.1 ± 2.0 nm for MFt and synomag, respectively. Moreover, from the DLS measurement a polydispersity index (PDI) of 0.11 (MFt) and 0.17 (PEGylated synomag) was derived. Overall, both particles show appropriate and comparable size homogeneity. Such size monodispersity is important to enable a robust determination of particle properties. It is also essential for any bio-technological or bio-medical application, since the MNPs’ response to a magnetic stimulus should be similar. The effective hydrodynamic diameter of < 50 nm is suitable for subcellular applications, since cytoplasmic non-specific interactions between a nanoprobe and the proteins/fibers in the cytoplasm dramatically increase above 50 nm, as previously reported [28]–[30].

**Figure 1.**
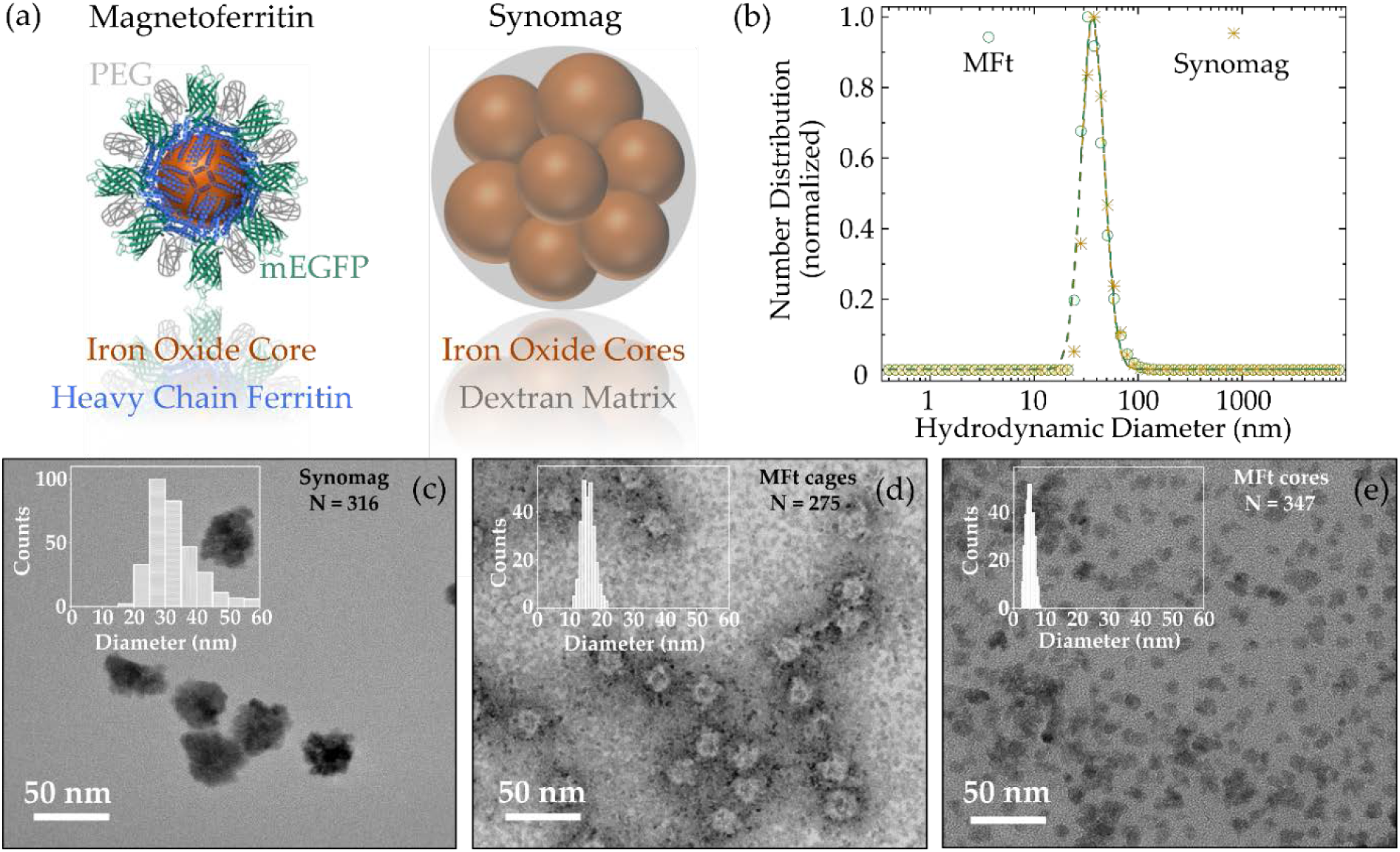
(a) Morphology sketches for magnetoferritin (MFt) [27] and synomag nanoparticles (NPs). (b) Number distribution for MFt (green) and PEGylated synomag (yellow) obtained by dynamic light scattering. Transmission electron microscopy images of (c) synomag, (d) MFt protein cages stained with uranyl acetate, and (e) MFt cores unstained. PEG = polyethylene glycol, mEGFP = monomeric enhanced green fluorescent protein, *D*_H_ = hydrodynamic diameter, *N* = number of evaluated NPs.

Transmission electron microscopy (TEM) images in Figure 1(c)-(e) were recorded to determine particle sizes and morphologies. TEM images of synomag MNPs (Figure 1(c)) confirm the ‘nanoflower’-like structure of the NPs with irregular surface texture. MFt protein cages are visualized using uranyl acetate (Figure 1(d)) and show the predicted uniform spherical structure. Note, that mEGFP and PEG on the cage surface are likely to give rise to the roughness increase of the outer cage. Magnetic cores after their synthesis into MFts are depicted in the unstained TEM image (Figure 1 (e)). Each TEM image is supplemented with a histogram and number of counted MNPs. The average diameter of synomag is 33.6 ± 7.6 nm, whereas dried MFt cores and cages are 5.1 ± 1.2 nm and 16.0 ± 2.0 nm, respectively (Table 1). In case of MFt, the sizes compare well with the protein crystalline structure with a cage inner diameter of 8 nm and an outer diameter of 12 nm [31]. Based on structural considerations mEGFP and PEG are predicted to add ~ 3 nm to the surface thickness [19]. Hence, the theoretically expected outer diameter of 18 nm compares well with our TEM results [32]. A comparison to the hydrodynamic diameters shows that synomag exhibits an 8.3 nm smaller diameter than its hydrodynamic diameter, which may be well explained by a ~ 4 nm hydrodynamic shell around the NPs [24]. MFt cages turn out to be about two times smaller than the hydrodynamic diameter. This discrepancy may either arise from larger iron oxide cores which form outside of the protein cages during the core synthesis step, or from remaining particle-particle interactions wherever the PEG passivation was insufficient to suppress all non-specific interactions. Finally, differences between the solid size and the size in the solution are expected from theoretical considerations, since the hydrodynamic size determination assumes diffusion of an ideal sphere, as reported before [33]–[35]. Subsequently, electrophoretic light scattering (ELS) measurements were performed to determine the NP surface potential, the so-called ζ-potential. For synomag and MFt the ζ-potential distribution exhibited a monomodal peak, confirming their surface homogeneity (data not shown) [35]. For both NPs the ζ-potential turned out to be weakly negative: -3.7 mV for MFt and -2.0 mV for synomag (Table 1). While both of these ζ-potential values deviate only slightly from zero, their negative charge may support colloidal stability, cellular uptake and cytoplasmic stealth properties, given the negative charge of many membrane and intracellular molecules [36], [37]. In addition, NPs of weak negative ζ-potential between -2 to -15 mV were recently shown to exhibit high intracellular mobility [29], [38].

**Table 1.** Physical parameters for MFt and PEGylated synomag: polydispersity index (PdI) from dynamic light scattering (DLS), *D*_TEM_ solid NP diameter from transmission electron microscopy (TEM), *D*_H_ hydrodynamic diameter from DLS, and ζ-potential from electrophoretic light scattering (ELS). MFt was stabilized in HEPES buffer (20 mM HEPES, 100 mM NaCl, pH 8.0), synomag in PBS (pH 7.4). Additional data can be found in Tables S1 and S2, Supplementary information.

| Sample | PdI | $D_H$,<br>nm | $D_{TEM}$,<br>nm | ζ-potential,<br>mV |
| --- | --- | --- | --- | --- |
| PEGylated<br>magnetoferritin | $0.11 \pm 0.01$ | $39.1 \pm 2.5$ | Cages: $16.0 \pm 2.0$<br>Cores: $5.1 \pm 1.2$ | -3.7 |
| PEGylated<br>Synomag | $0.17 \pm 0.03$ | $39.1 \pm 2.0$ | $33.6 \pm 7.6$ | -2.0 |

A major challenge in realizing a remote and efficient manipulation of MNPs in the cell environment is to adjust the NPs magnetic properties, such as their magnetization and magnetic anisotropy, to the external magnetic fields. To characterize the magnetic response of the herein presented NPs, conventional magnetic field- and temperature-dependent vibrating sample magnetometry (VSM) was conducted. Figure 2(a)-(b) shows magnetization curves for MFt and synomag. At *T* = 300 K both samples exhibit superparamagnetic behavior and a saturation magnetization of 42 Am^2^/kg (MFt) and 69 Am^2^/kg (synomag). The insets of Figure 2(a)-(b) present the low field region. The superparamagnetic behavior is essential to reduce magnetic dipole-dipole interactions between the particles and hence their propensity to form chains. Exhibiting this property at physiological temperatures is further important to enable a manipulation of each particle individually without any application of torque to the NPs. At *T* = 5 K, both samples exhibit a hysteresis loop with coercive fields of 31 mT and 8 mT for MFt and synomag, respectively. This is a typical signature of the ferrimagnetic nature of the respective Fe oxides. The saturation magnetization at *T* = 5 K is 56 Am^2^/kg for MFt and 81 Am^2^/kg for synomag which are both higher than the values at *T* = 300 K, as expected, but lower than the volume magnetizations of bulk magnetite (92 Am^2^/kg) and maghemite (82 Am^2^/kg) [39].

**Figure 2.**
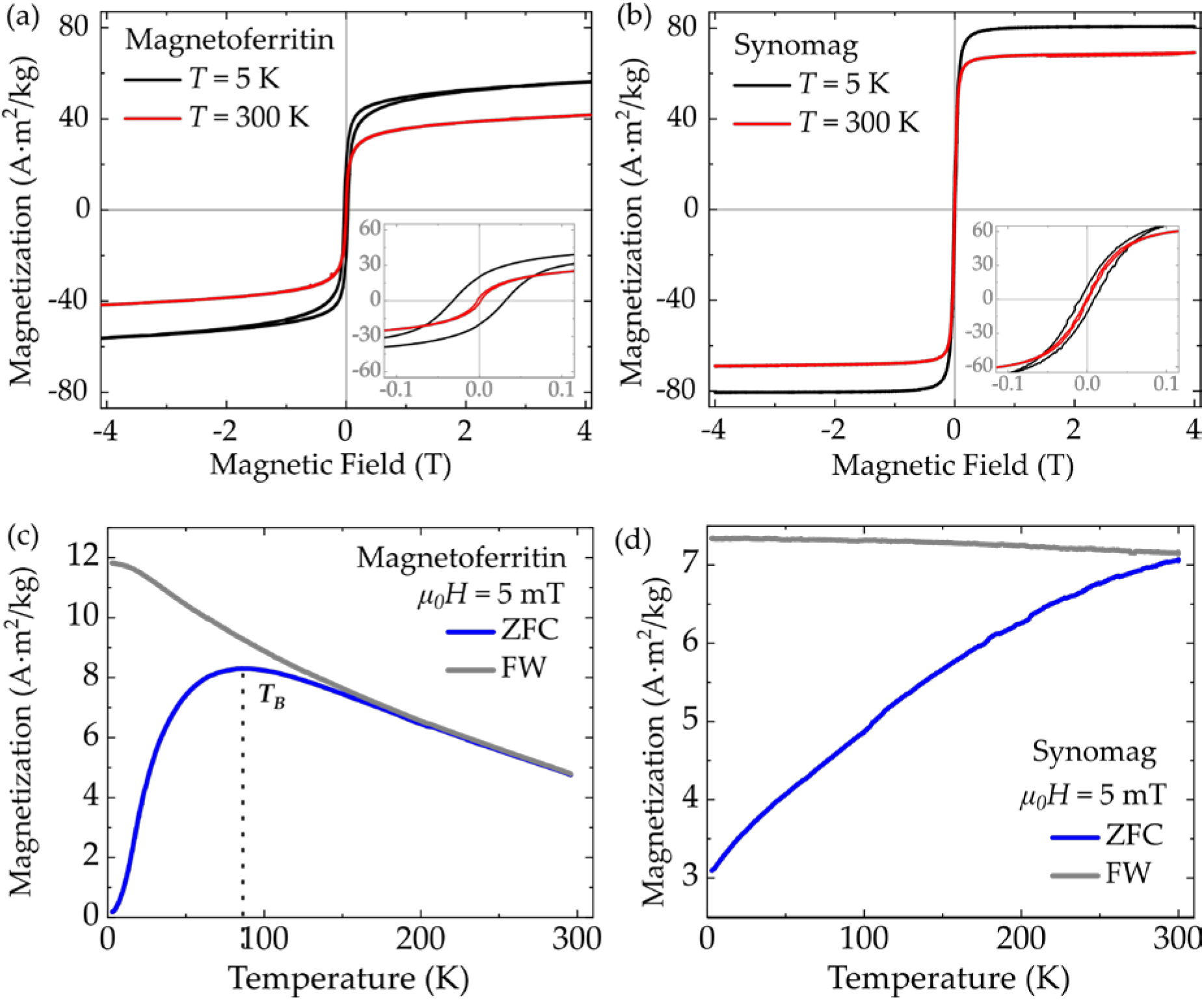
Magnetization as function of magnetic field for (a) magnetoferritin and (b) synomag nanoparticles. ZFC/FW for (c) magnetoferritin and (d) synomag measured in 5 mT. *T_B_* = characteristic blocking temperature, ZFC = zero field cooling, FW = field warming.

Figure 2(c)-(d) shows zero-field-cooling/field-warming (ZFC/FW) experiments. MFt exhibits a broad maximum centered at 86 K, which indicates the effective blocking temperature *T_B_*, and an irreversibility point at about 150 K. The MFt ZFC/FW maximum at 86 K found in this work is significantly larger than the *T*_B_ reported in [40] (*T_B_* = 40 K) and in [41] (*T_B_* = 13 K). This shift in *T*_B_ may result from reduced distances and dipole-dipole interactions between cores in the powdered samples after the drying process. Here, in order to minimize paramagnetic contributions to the VSM data, the salt in the medium was strongly diluted, which in turn may affect protein cage stability. Other contributions may arise from interactions between cores synthesized outside of the cages. Since for VSM measurements with an averaging time of 1 s and blocking temperature *T_B_* the relation 21·*k*_B_*T*_B_ ≈ *K*_eff_*V* holds [41], the effective magnetic anisotropy value *K*_eff_ can be estimated with the particle volume V. Using the mean particle diameter of 5-6 nm, we obtain *K_eff_* = 2-4 · 10^5^ J/m^3^. This value is reasonable for Fe oxide MNP of small sizes, exhibiting a large surface to volume ratio. Here, magnetic moments at the surface of a MNP experience broken symmetry and crystal deterioration leading to a larger surface anisotropy compared to volumetric magnetite [39], [42]. Such increasing effective magnetic anisotropies are typically observed with reduced MNP sizes [43]–[45].

For synomag, experimental magnetic field and temperature dependencies of the magnetization obtained in this work complement the characterization previously reported in [25]. We obtain a broad distribution of blocking temperatures with an anticipated center slightly above physiological temperature (see Figure 2(d)). This indicates that synomag MNPs are designed to effectuate maximal hysteretic losses in radio-frequency alternating magnetic fields and thus exhibit good heating capabilities for magnetic particle hyperthermia [17], [25]. Taken together both MNP systems possess the key magnetic properties (superparamagnetism, sufficient magnetic response) necessary to realize a remote manipulation in external magnetic fields.

Despite protective coatings, iron oxide MNPs are still cautiously applied in nanomedicine due to potential side effects they could implicate after being injected into living tissue or cells [1], [46]. Here, HeLa WT cells were used to test the cell viability with the CellTiter-Blue assay on both MNP systems within the concentration range of 0.1-4.0 mg/ml. MNPs were transferred into the cells via a standard incubation protocol. As controls, untreated cells in DMEM and cells treated with MNP buffers were used. As an additional control, cells were killed by addition of 0.1% Triton-X for 30 min before imaging was started. As shown in Figure 3, traceable harmful effects for both samples appeared only at concentrations above 1.0 mg/ml. For ferritin cages (Figure 3(a)), a small reduction in viability was also detected in the control measurement with HEPES buffer (20 mM HEPES, 100 mM NaCl, pH 8.0). Similarly, a viability drop for synomag only occurred at concentrations above 1.0 mg/ml. Addition of PBS did not affect the cell viability. In conclusion, both MNPs can be safely brought into the cell environment at concentrations up to 2.0 mg/ml.

**Figure 3.**
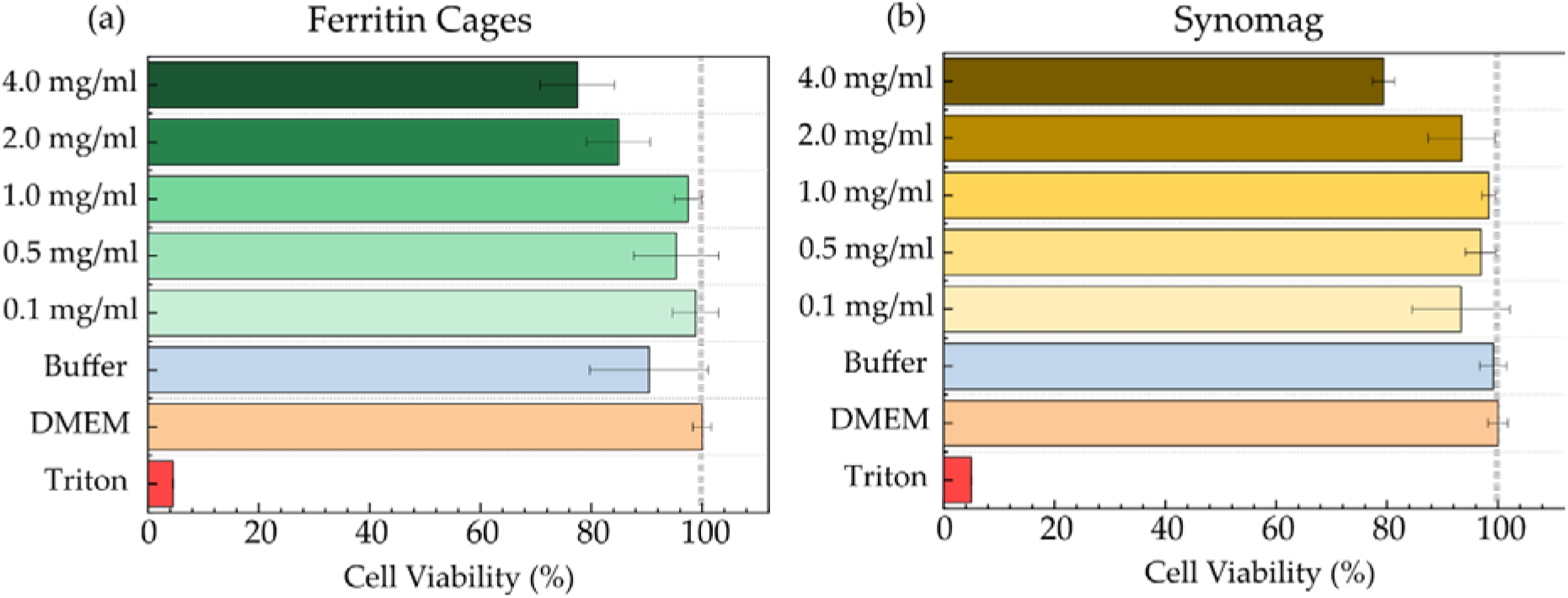
Cell viability Assay (CellTiter-Blue from Promega) measured after 24 hours in HeLa WT cells for (a) ferritin cages and (b) synomag. Data is normalized to DMEM as positive control with untreated cells. 0.1% Triton leads to cell death and is therefore used as negative control. The buffer control determines the effect of plain buffer on cell viability. The highest volume of MNP solution that was added during MNP addition was used here. Buffers were 20mM HEPES, 100 mM NaCl (pH 8.0) for ferritin and PBS (pH 7.4) for synomag.

Hence, for subsequent particle incubation experiments the upper concentration limit was set to 1.0 mg/ml (equals to 6.4 μM for MFt and 18.3 pM for synomag) to rule out any negative MNP related effect. Previous studies investigating the effects of iron oxide NPs on cell viability reported similar results, where multi-core iron oxide NPs passivated with citrate of up to 5 mM showed little negative effects on cells [47]. Also PEGylation and passivation was reported to reduce cellular damage upon uptake [48], [49], supporting our particle passivation strategy.

In order to characterize the process of particle incubation and uptake further and in view of subsequent magnetic manipulation, where an efficient particle delivery into the cell is desirable, we evaluated the intracellular NP amount via image data analysis of their fluorescence. NP uptake by HeLa WT cells was monitored at distinct incubation times of 6 and 24 hours to probe different long-time temporal effects as well as for concentrations between 0.1-1.0 mg/ml. In this study, PEGylated Ft cages without a magnetic core were used, since magnetoferritin is freshly produced only in small amounts and since the magnetic properties are not relevant for this assay. Care was taken to record the cell samples at identical imaging settings, in order to not bias the detected fluorescence intensities. Obtained data was evaluated using a custom-written Matlab algorithm. Therein, the intensity per pixel inside each cell as a function of NP concentration was extracted and normalized to the amount of cells found (see Figure 4(a) for Ft cages and Figure 4(d) for synomag MNPs). Ft cages demonstrated a substantially larger cellular uptake compared to synomag with an increase of the MNP concentration inside the cell if incubated with higher MNP concentration. Evaluation of the uptake for different concentrations after 6 h of incubation shows only a slight increase in the recorded intensity of both particles even at the highest concentration of 1 mg/ml (Figure 4(d)). After 24 h a higher amount of MNPs is taken up and this is increased with increasing particle concentration in the medium. Figures 4(b),(e) show phase contrast images of the cells and Figures 4(c),(f) are the corresponding fluorescence images indicating the MNP distribution after 24 h of incubation. In Figure 4(c) Ft cages show a cloud-like accumulation around the nucleus. This is a typical signature of particles which are taken up via the endosomal pathway. Here, after initial encapsulation within the endosome, the cargo is transported towards the endoplasmic reticulum or Golgi apparatus, where it is further processed. In contrast, synomag MNPs are more sparsely distributed, accumulating around the nucleus as well as in spots across the cell (see Figure 4(f)). Interestingly, according to Figure 4(a) and (d), Ft NPs are taken up more efficiently compared to synomag MNPs. This may be attributed to the twofold difference in their solid diameters. Previous studies already reported a positive effect of a decreasing particle size on cellular uptake [50]. The slight negative ζ-potential (see Table S1 and S2) measured for both NPs may also enhance particle uptake, since most intracellular proteins and lipids are negatively charged. The negative surface charge can further protect NPs against non-specific interactions inside the cell, preventing their entrapment [29]. For our MNPs, the stealth properties were further enhanced by modifying the NPs surface using PEG as passivating agent, as previously suggested [34], [38], [50].

**Figure 4.**
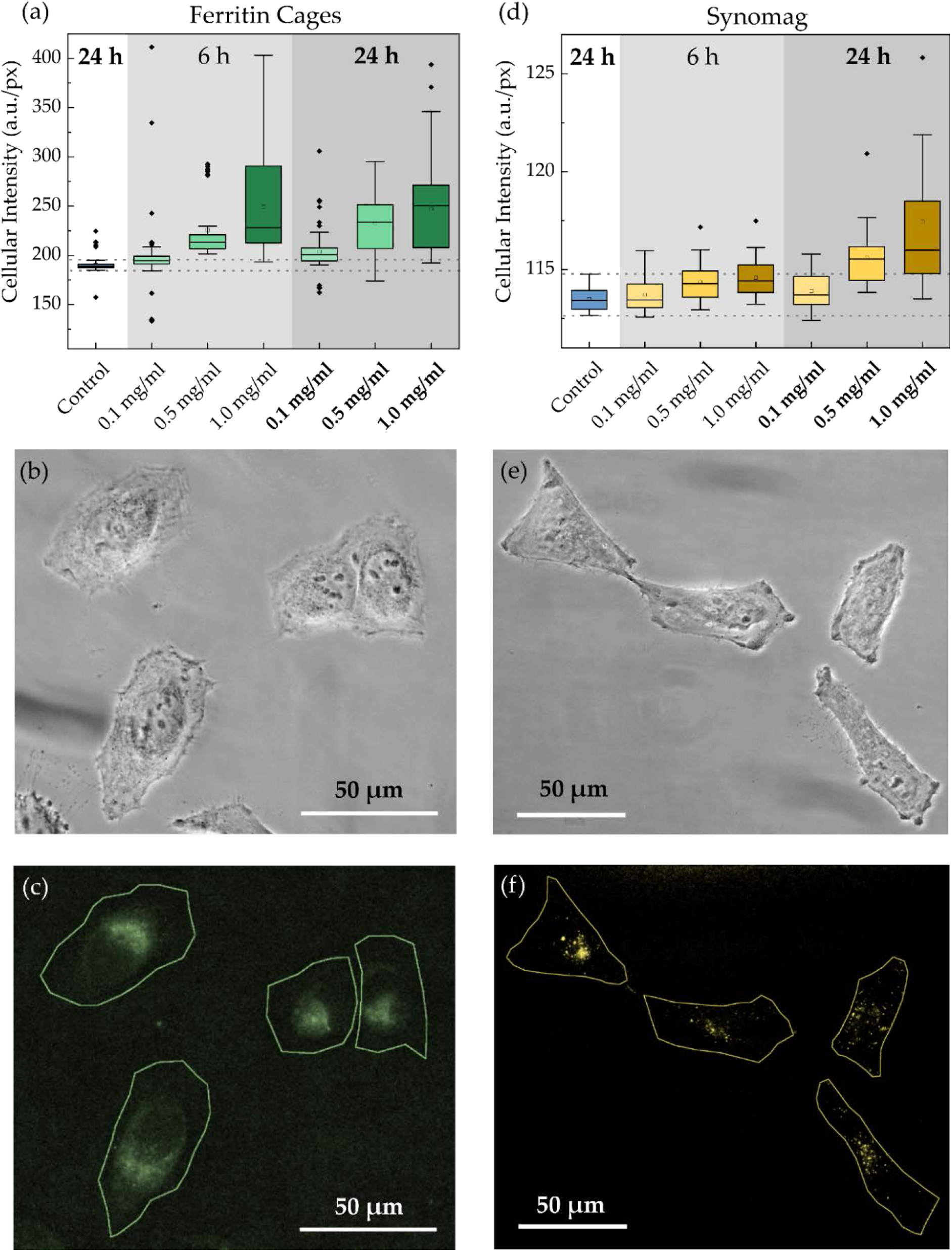
HeLa WT cellular uptake of (a) ferritin cages and (d) synomag nanoparticles (NH_2_ surface) as function of concentration. Intensity is given as intensity per pixel inside the cell. Particle systems were transferred into the cells via incubation for 6 or 24 hours. (b)-(c) Phase contrast image and corresponding fluorescent image with cell outlines recorded using 470/525 nm filter for ferritin cages. (e)-(f) Phase contrast image and corresponding fluorescent image with cell outlines recorded using 545/620 nm filter for synomag NPs.

**Figure 5.**
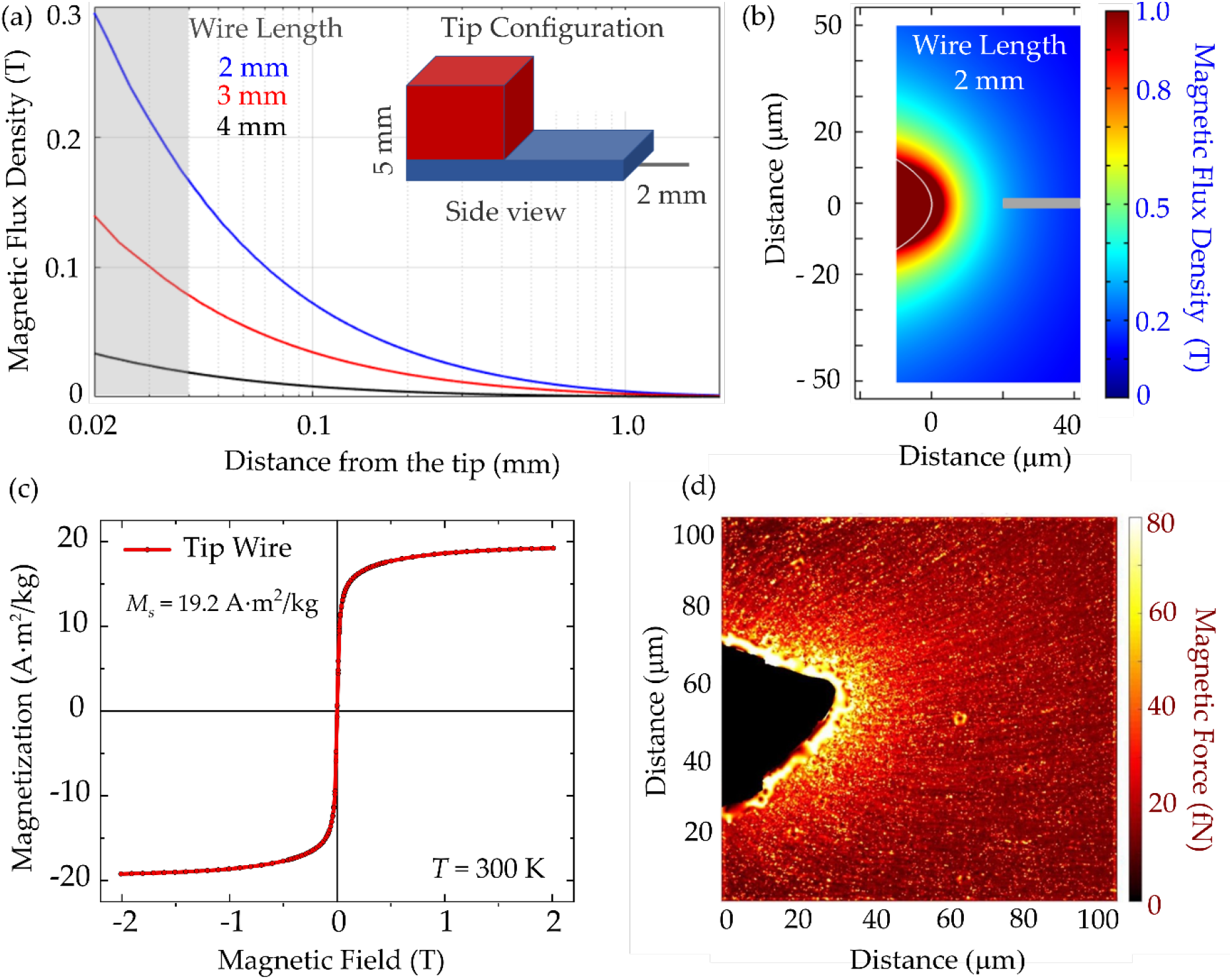
Magnetic tip for remote manipulation. Simulated magnetic flux density (a) for a tip wire extending 2, 3, or 4 mm from the magnet as a function of distance from the tip, and (b) as 2D projection from the top with 2 mm long tip wire. Grey line in (b) indicates the position from where the data in (a) - highlighted by the grey area - was extracted for the 2 mm tip wire. Inset in (a) shows a sketch of the tip configuration consisting of two magnets (red, blue) and a tip wire of 2 mm attached to the magnets. (c) Magnetization curves of the tip wire measured at *T* = 300 K. (d) Magnetic force measured via nanoparticle attraction to the magnetic tip with 2 mm tip wire. The line texture resulted from the tracked particles.

After successful uptake, we then sought to realize the remote and localized manipulation of MNPs inside cells using a magnetic tip. To this end, first the magnetic tip and the generated magnetic flux density gradient had to be characterized. The magnetic tip was set up as a combination of two permanent magnets and a string wire was attached to the lower magnet (Figure 5(a), inset). The string wire was made of spring steel and pulled into a tip of parabolic-like shape. To characterize the magnetic flux density generated by the magnetic tip, 3D finite element modeling as well as an *in vitro* calibration assay were performed. In case of finite element modeling, the magnetic flux density was simulated for 2, 3, and 4 mm lengths of the tip wire and plotted against the distance from the tip (see Figure 5(a)). Non-surprisingly, the 2 mm long tip exhibits the highest magnetic flux densities at small distances that are critical for remote manipulation. Moreover, the strong dependency of the absolute magnetic flux density values indicates how important an accurate choice of the wire length is. The grey area in Figure 5(a) corresponds to the grey line in Figure 5(b), where the simulated magnetic flux density around the magnetic tip is shown as 2D projection from the top. Figure 5(c) depicts the magnetization curve of the tip material with a saturation magnetization of 19.2 Am^2^/kg and a coercive field of 1.2 mT. Figure 5(d) shows the result of an *in vitro* assay, which was introduced to measure the magnetic force generated by the magnetic tip over 100 μm distance. This is the relevant scale for particle manipulation experiments in single cells.

To realize this assay, precise positioning of the tip inside the sample was achieved using an InjectMan 4 micromanipulator, where the three spatial axes as well as the angle of the magnetic tip can be adjusted with 20 nm spatial precision. The tip was positioned in a mixture of 85% glycerin and 15% distilled water, to mimic intracellular viscosity conditions. As magnetic nanoparticle probes, Nanomag-D with 250 nm hydrodynamic diameter (micromod GmbH, Rostock, Germany) were used. These enabled individual detection and reliable tracking with phase contrast microscopy. MNPs within the glycerin/water mixture were attracted toward the tip and the trajectories of several 100 particles were recorded. Thereafter, the Fiji plugin TrackMate and self-written routines were used to trace the particle trajectories and to calculate the velocity vectors along the particle trajectory [51].

Using Stokes’ law

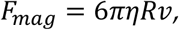

the magnetic force generated by the tip was calculated, which typically lay within the range of 10-100 fN. Here, *η* is the dynamic viscosity of the medium as determined in [29], *R* is the MNP radius, and *υ* is the velocity of MNPs extracted from the single particle trajectories.

Having established the above prerequisites for the magnetic manipulation approach, we probed MNP transfer into HeLa WT cells and their subsequent intracellular mobility via two approaches: MNP injection using a microneedle (Figure 6) and MNP incubation (Figure 7). In both cases, MNP localization was recorded via fluorescence microscopy in the presence of the magnetic tip for about 30 min followed by recordings in the magnetic tip’s absence. Fluorescence intensity distribution inside a single cell was monitored and plotted as a function of time for regions within the magnetic tip’s sphere of action. The mean intensity *I* for this region as a function of time was fitted by an exponential fit function to obtain the characteristic accumulation *τ*_acc_ and relaxation *τ*_rel_ times:

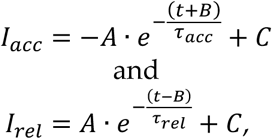

**Figure 6.**
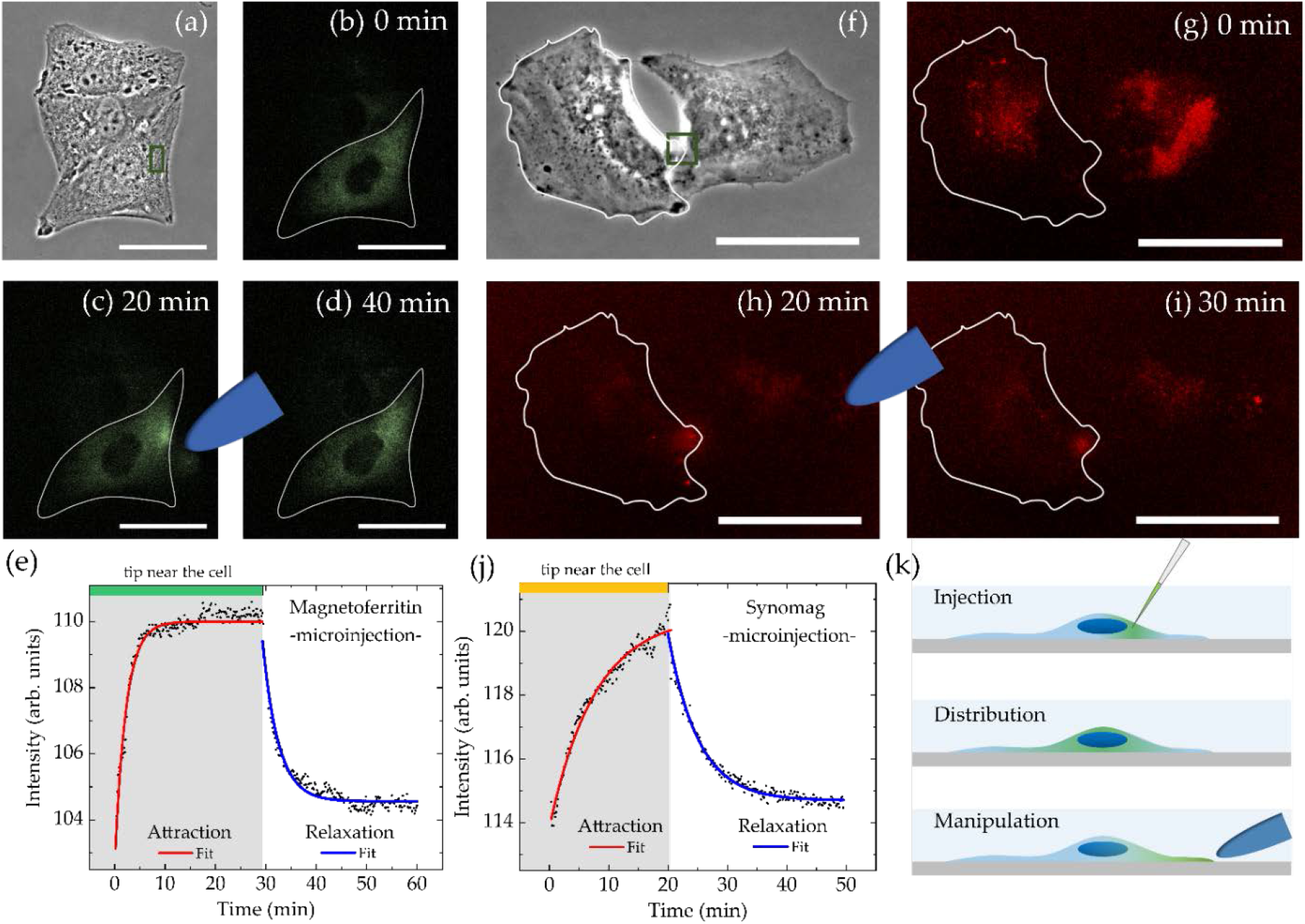
Spatial manipulation of (a)-(d) magnetoferritin and (f)-(i) synomag nanoparticles inside living HeLa WT cells after microinjection recorded over 40 min. Scale bars are 50 μm. (e),(j) show corresponding mean intensities over time with fit functions to characterize nanoparticle accumulation near the magnetic tip (green box in (a) and (f)) and relaxation after tip removal. Distance between magnetic tip and analyzed region was (MFt) 6 μm or (synomag) 75 μm respectively. (k) illustrates the experimental setup for microinjection.

**Figure 7.**
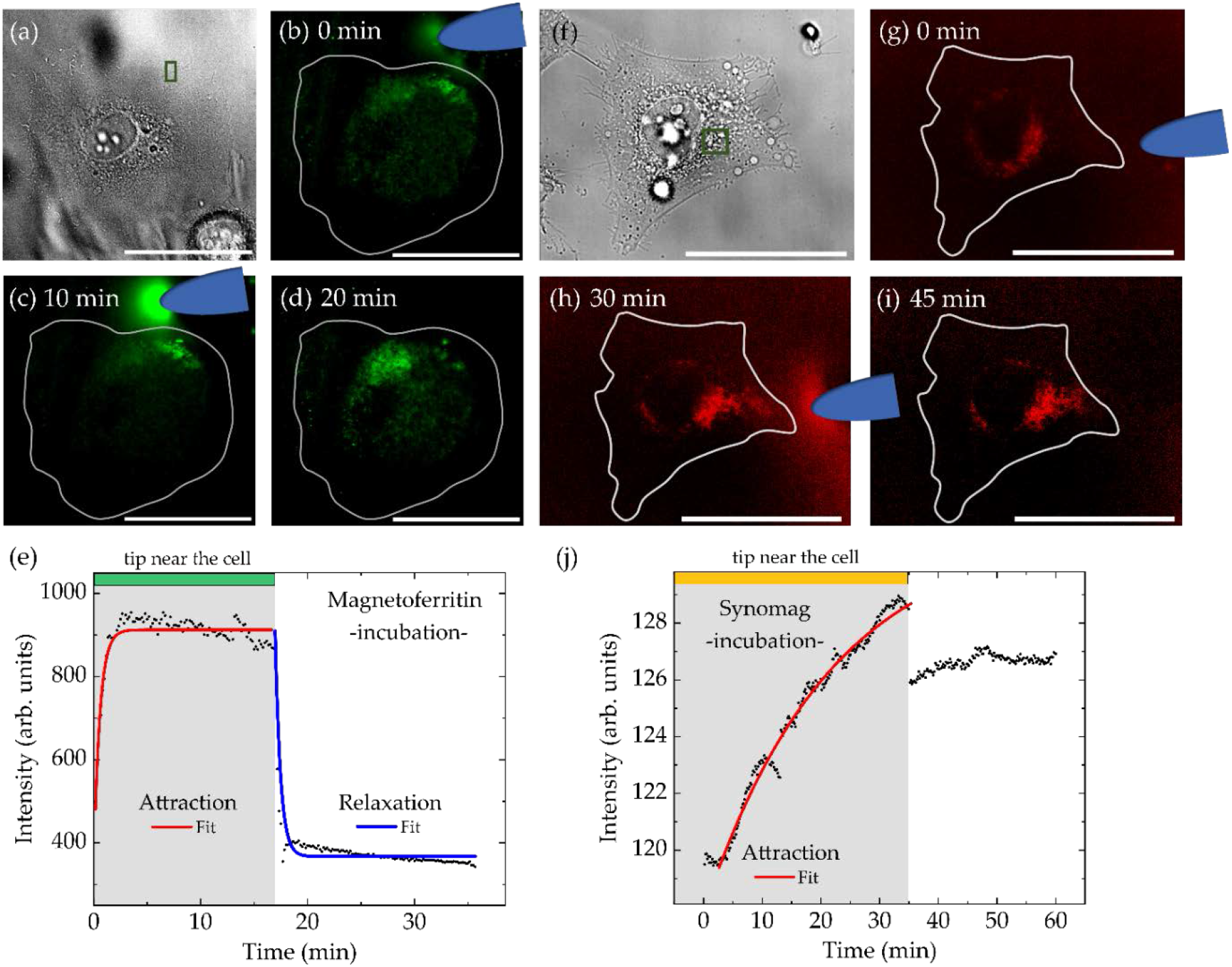
Spatial manipulation of (a)-(e) magnetoferritin and (f)-(j) synomag nanoparticles inside the single HeLa WT cell after 5 and 24 h incubation, respectively. Scale bars are 50 μm. (e),(j) show corre-sponding mean intensity over time along with the fit to characterize accumulation time near the magnetic tip (green box in (a) and (f)) and relaxation time after the tip removal. Distance between magnetic tip and analyzed region (green box in (a) and (f)) was (MFt) 10 μm or (synomag) 30 μm respectively.

where *A* is a fit parameter reflecting the intensity plateau of the exponential function on long time scales, *B* represents the temporal off-set of the starting point of attraction/relaxation, *C* stands for the intensity offset, and *τ*_acc_ and *τ*_rel_ are characteristic accumulation and relaxation times, respectively.

Injection was performed using a microinjector (FemtoJet 4i) in combination with a micromanipulator (InjectMan 4). For particle delivery, a microinjection capillary (Femtotip II) with an inner diameter of 500 nm was used. Dispersed particles should effortlessly pass through this capillary underlining the importance of using a highly homogeneous and non-reactive MNP sample. MFt with PEGylated surface was readily injected. Synomag with a NH_2_ coating was tested initially, but experienced aggregation inside the injection needle, which irreversibly blocked it. Since surface passivated MNPs are expected to exhibit reduced parti-cle-particle interactions and to be less prone to aggregation, the PEGylation protocol for MFt was also conducted for synomag NPs. The effective hydrodynamic diameter after PEGylation changed from 41.9 ± 0.5 to 39.1 ± 2.0 (Table S2, Supplementary Information) and was hence similar to the one of MFt (Figure 1). This PEG passivation indeed facilitated synomag microinjection.

Injected MFt (see Figure 6(a)-(d)) were instantaneously homogeneously distributed after microinjection into the cell. It was possible to both attract and accumulate particles using a magnetic tip. Once the magnetic tip was removed, MFts returned to the initial homogeneous distribution within minutes. Figure 6(e) shows the relatively fast kinetics with a characteristic accumulation time *τ*_acc_ of 2.2 ± 0.1 min and a relaxation time *τ*_rel_ of 3.1 ± 0.1 min. Very similar values were reported in a previous study, where PEGylated core-shell maghemite-silica particles with diameters of about 40 nm were accumulated within *τ*_acc_ of ~ 1 min, whereas the relaxation times *τ*_rel_ were ~ 10 min with a large distribution between 2 and 35 min [28].

In contrast, obtaining a homogeneous particle distribution after synomag injection appeared to be more difficult than with MFt. An example of microinjection of PEGylated synomag into HeLa WT is given in Figure 6(f)-(i). Synomag MNPs exhibited heterogeneous particle distribution inside cells and despite the PEG passivation were still prone to entrapment in the cytoplasm. Nevertheless, in some cases, MNPs could be attracted via the magnetic tip. After magnetic tip removal the relaxation dynamics were recorded and exhibited significant loss in mobility for synomag compared to MFt. Attraction and relaxation times for the case shown in Figure 6 increased to *τ*_acc_ = 5.1 ± 0.1 min and *τ*_rel_ = 7.7 ± 0.2 min. Intriguingly, synomag attraction towards the magnetic tip could be monitored for distances up to 100 μm. This can be attributed to the higher saturation magnetization and magnetic moment (Figure 2(b)), corresponding to higher forces of synomag compared to MFt. Thus, higher forces can develop and attract particles across the cell over large distances, despite increasing non-specific interactions. Qualitatively, successful particle attraction was observed in around 10 % of the analyzed cells, of which 50 % exhibited a relaxation of particle distributions after tip removal.

In case of magnetic manipulation after particle incubation, we considered the results of our particle uptake study as presented in Figure 4. Choosing incubation conditions of 0.5 mg/ml of MFt in the extracellular medium over 24 h led to the expected high particle uptake. However, MFts could not be attracted by a magnetic tip, not even over distances less than 10 μm. Most probably this was due to increasing intracellular interactions and particle recognition by the cell’s autophagy machinery on long time scales. Therefore, shorter incubation times of 1, 2, 3, 4, and 5 h with 0.5 mg/ml of MFt in the extracellular medium were tested. Cells were washed and upon observation under the microscope, those cells exhibiting sufficient MFt signal were monitored for 1 h while approaching and removing a magnetic tip.

Figure 7(a)-(d) shows a cell after incubation with MFt for 5 h. Here, prior to magnetic manipulation, MFts were evenly distributed across the cell. Magnetic particles were attracted with a characteristic time of *τ*_acc_= 0.56 ± 0.04 min and redistribution in absence of the magnetic tip occurred with a characteristic time of *τ*_rel_ = 0.53 ± 0.03 min. These values are distinctively smaller than values from injection studies. Changes in the attraction kinetics may in general also be attributed to variable distances between the magnetic tip and the MNPs, since the magnetic attraction force scales proportional to 1/r^2^, where r is the distance. However, in this case both, attraction and relaxation dynamics evolved on fast timescales, suggesting that MFts did not enter the highly viscous cytoplasm of the cell. Instead, after the short incubation time used, MFt may be attached on the outer side of the cell plasma membrane, wherefore particles could be more easily attracted towards the tip. In conclusion, manipulation of MFt after incubation was difficult to realize as in most of the cases MFt remained immobile. Thus, magnetic forces in the fN range will only attract the MNPs when the applied force surpasses any intracellular interaction. Note, that for longer measurement times the magnetic tip became increasingly fluorescent due to freely moving MFt particles in the extracellular medium. These were present despite several washing steps prior to imaging. While this was disturbing during imaging, the MNP attraction by the tip provided evidence that the particles were still magnetic after 24 h incubation (see Figure 7 (c)).

In Figure 7(f)-(j) an example of incubation and manipulation of PEGylated synomag is shown. In contrast to MFt, synomag manipulation was possible even after 24 h of incubation. The cell shown in Figure 7(f) was incubated with a concentration of 0.5 mg/ml for 24 h, washed and then analyzed under the microscope. Although being mostly located around the nucleus, PEGylated synomag MNPs were still attracted by the magnetic tip. The accumulation time *τ*_acc_ amounted to 21.8 ± 0.3 min, and was significantly higher than for MFt and synomag after MNP injection. As discussed in the context of Figure 4, this may be attributed to an entrapment of synomag inside cellular vesicles during particle uptake. In line with this interpretation, no intensity relaxation and particle redistribution after magnetic tip removal was observed (Figure 7(j)). Instead, similarly to Figure 7(c) the magnetic tip gradually accumulated remaining MNPs from the extracellular medium and showed a fluorescent signal, confirming their magnetic response even after 24 h of incubation. However, as a result of this attraction, a distinct drop of intensity was observed upon tip removal, after which intensities remained constant (see Figure 7(j)). Accordingly, the fit for the relaxation time determination in this case became obsolete. Based on the aforementioned results concerning synomag, we hence conclude that the higher magnetic forces exerted by PEGylated synomag - in contrast to MFt - can exceed the non-specific interaction forces. The particles as well as any coupled molecule may then be pulled across the cell.

The possibility to manipulate MNPs after incubation and uptake by cells is of particular interest, since MNP incubation affects multiple cells simultaneously and is considered a multicellular approach. Understanding MNP incubation and uptake is further useful in magnetic cell sorting studies, as suggested by Massner et al. [41]. In addition, the possibility to transfer localized forces to the cell is important whenever mechanical forces influence a cellular process. For example Tseng et al. showed that the cell orientation during cell division is susceptible to MNP mediated force application [52] and Seo et al. [9] demonstrated how mechanosensitive receptors on the cell surface influence cell expression. Here, we have provided a proof of principle, under which conditions such force stimulus can be applied.

## 4. Conclusions

In this work, we compared two new classes of MNPs, semisynthetic single core magnetoferritin (MFt) and multicore synomag MNPs, in view of future magnetic manipulation studies in single cells. Both particles fulfilled the prerequisite of size monodispersity and high biocompatibility up to MNP concentrations of 2 mg/ml. Efficient intracellular transfer of MNPs was realized by simple MNP incubation, which is advantageous for nanomedicine applications compared to other transfer methods. Higher transfer efficiencies in case of MFts compared to synomag were attributed to the smaller size and the slightly more negative ζ-potential. The difference in magnetic core size further gave rise to varying forces in in vitro experiments, with the larger magnetic moments of synomag developing higher magnetic force responses than MFts. These MNP characteristics were ultimately tested in two subcellular manipulation approaches: 1) particle microinjection, where free particle motion in the cell cytoplasm and reversible particle redistribution via a magnetic tip was realized. Here, MFts exhibited higher mobility and reversible attraction-relaxation kinetics compared to synomag NPs. The second manipulation approach was based on 2) particle incubation and uptake by cells. Here, most NPs accumulated at cell organelles and exhibited reduced mobility. Magnetic field gradients probed the NP mechanical force response, which successfully led to synomag attraction across the cell, whereas MFts remained immobile. Hence, MFts are more suitable for non-invasive spatial manipulation approaches in cells, whereas synomag are well utilized to mediate nanoscale forces. The experimental assays described and the values reported for MNPs provide benchmarks for future magnetic manipulation experiments, whenever a spatial or mechanical manipulation of a biological process is envisaged.

## Supplementary Materials

The following are available online at www.mdpi.com/xxx/s1, Figure S1: SDS-PAGE analysis of purification steps of ferritin shells; Table S1: Hydrodynamic size, polydispersity index and ζ-potential for subsequent steps of magnetoferritin synthesis; Table S2: Hydrodynamic size, polydispersity index and ζ-potential for synomag nanoparticles with three surface modifications.

## Author Contributions

Conceptualization, C.M. and I.N.; methodology, C.M., I.N., A.N., U.W., and N.B.; software, A.N. and J.S.B.; investigation, I.N., A.N., J.S.B., U.W., M.R.S. and M.O.; resources, C.M., M.K., and M.F.; writing—original draft preparation, I.N.; writing—review and editing, C.M., I.N., A.N. and U.W.; visualization, I.N. and A.N.; supervision, C.M. and I.N.; project administration, I.N.; funding acquisition, C.M., M.K., and M.F. All authors have read and agreed to the published version of the manuscript.

## Funding

C. M. acknowledges financial support of the Deutsche Forschungsgemeinschaft (DFG) through SFB1208 ‘Identity and Dynamics of Membrane Systems’ (A12). C.M., I.N. and A.N. acknowledge financial support via the ‘Freigeist fellowship’ of VolkswagenFoundation. This work was performed on synomag particles purchased from Micromod Partikeltechnologie GmbH with financial support from Fonds der Chemischen Industrie. The authors acknowledge the DFG and the State of North Rhine-Westphalia for funding the cryo-TEM (INST 208/749-1 FUGG).

## Data Availability Statement

We can provide original data upon request.

## Acknowledgments

We acknowledge Prof. Dr. C. Seidel and Prof. Dr. L. Schmitt and their groups for sharing their biochemistry equipment and expert knowledge on its handling. We thank the group of M. Coppey and B. Hajj at the Laboratoire Physico-Chimie, Institut Curie, Paris, France for providing the mEGFP::HCF plasmid. C.M. acknowledges Maxime Dahan^†^, Institut Curie, for introduction into the topic. We thank Daniel Kuckla for microscopy support, optimization of magnetic core loading setup and useful comments to improve the manuscript.

## Conflicts of Interest

The authors declare no conflict of interest.

## Notes

### Competing Interest Statement

The authors have declared no competing interest.

