## Supplementary Information for "Magnetic Nanoprobes for Spatio-Mechanical Manipulation in Single Cells"

[1] Experimental Medical Physics, Heinrich-Heine University Düsseldorf, 40225 Düsseldorf, Germany
[2] Colloids and Nanooptics, Heinrich-Heine University Düsseldorf, 40225 Düsseldorf, Germany
[3] present address: Peter Grünberg Institute, Electronic Properties (PGI-6), Forschungszentrum Jülich, 52425 Jülich, Germany
[4] Faculty of Physics and Center for Nanointegration (CENIDE), University of Duisburg-Essen, 47057 Duisburg, Germany
† contributed equally

### Ferritin Purification

Sodium dodecyl sulphate–polyacrylamide gel electrophoresis (SDS-PAGE, 12%) carried out to confirm ferritin quality after each purification step can be seen in **Figure S1**. A standard Coomassie blue staining protocol was used. Theoretical molecular weight (MW) for mEGFP::HCF monomer is 48.7 kDa. This expected band is clearly observed in all loaded gel pockets indicating the presence of the desired mEGFP::HCF. Bright green color of the solution further confirms the correct expression of the complex containing mEGFP. Sephacryl S400 16/60 size exclusion column (SEC) equilibrated in buffer (20mM HEPES pH 8,0, 100mM NaCl, pH 8.0) was used for the final purification step.

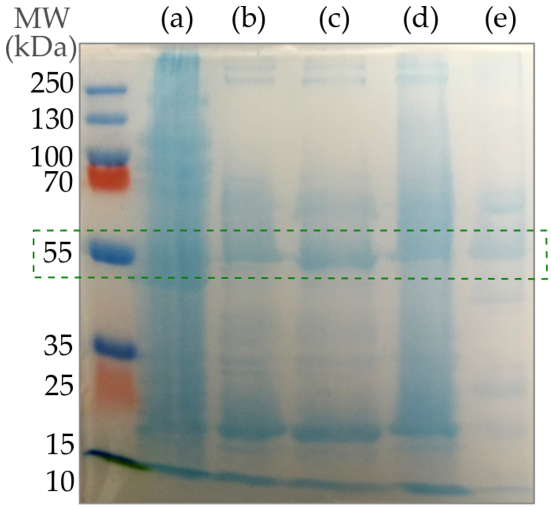


**Figure S1.** SDS-PAGE analysis of purification steps of ferritin shells. (a) directly after disrupting *E.coli* walls, (b) after heat denaturation at 70°C, (c) after 30% ammonium sulfate precipitation, (d) after 70% ammonium precipitation, (e) final product before loading on a size exclusion column (SEC).

**Table S1.** Hydrodynamic size $D_H$, polydispersity index (PdI), and ζ-potential for subsequent steps of magnetoferritin synthesis. Concentration c is presented after the filtering step (0.2 μm cutoff, PTFE). Stabilizing buffer (20 mM HEPES, 100 mM NaCl, pH 8.0) was used for all listed measurements. Conductivity of the medium was at 11 mS/cm for all samples while it was 0.1 mS/cm for milliQ water.

| **Sample** | **$D_H$, nm** | **PdI** | **ζ-potential, mV** |
|---|---|---|---|
| Non-PEGylated Ferritin Cages c = 0.5 mg/ml | 15.3 ± 1.0 | 0.25 ± 0.00 | -4.8 ± 0.7 |
| PEGylated Ferritin Cages c = 1.0 mg/ml | 20.5 ± 3.5 | 0.21 ± 0.01 | -3.5 ± 0.7 |
| PEGylated Magnetoferritin c = 0.7 mg/ml | 39.1 ± 2.5 | 0.11 ± 0.01 | -3.7 ± 1.2 |

**Table S2.** Hydrodynamic size $D_H$, polydispersity index (PdI), and ζ-potential for synomag nanoparticles with three different surface modifications. Concentration prior to filtering (0.2 µm cutoff, PTFE) was 1.0 mg/ml. Stabilizing buffer PBS (pH 7.4) was used for all listed measurements. Deviations from the ζ-potential are ascribed to the unpronounced phase plot prohibiting improvements of the final result disregarding the subruns increase.

| Synomag | $D_H$, nm | PdI | ζ-potential, mV |
|---|---|---|---|
| plain | 48.1 ± 1.5 | 0.11 ± 0.01 | -3.8 ± 0.6 |
| $NH_2$ | 41.9 ± 0.5 | 0.05 ± 0.01 | -1.2 ± 1.2 |
| $NH_2$ - $PEG_{2000}$ | 39.1 ± 2.0 | 0.17 ± 0.03 | -2.0 ± 2.3 |
